# Analysis of Divergent Gene Expression between HPV+ and HPV- Head and Neck Squamous Cell Carcinoma Patients

**DOI:** 10.1101/2025.04.14.648798

**Authors:** Kasturika Shankar, Sarah E. Walker

## Abstract

Human Papillomavirus (HPV) is a non-enveloped virus with a circular double-stranded DNA genome. It is one of the most common sexually transmitted infections, with high-risk types such as HPV-16 and HPV-18 linked to anogenital and head and neck squamous cell carcinomas (HNSCC). HNSCC includes cancers of the oral cavity, pharynx, larynx, and related regions, caused by carcinogens or persistent viral infections. HPV-positive (HPV+) HNSCC cases are more prevalent in Western countries and exhibit better prognosis and treatment response compared to HPV-negative (HPV-) cases. These differences suggest distinct fundamental differences between each subtype.

This study analyzed RNA-seq data from the PanCancer Atlas 2018 dataset to investigate molecular distinctions between HPV+ and HPV-HNSCC. Using dimensionality reduction techniques such as Principal Component Analysis (PCA) and Uniform Manifold Approximation and Projection (UMAP), a clear clustering of HPV+ cases was observed, suggesting a unique gene expression profile. HPV+ tumors exhibited upregulation of genes involved in nucleic acid processing and downregulation of genes associated with apoptosis and epidermis development. These findings underscore the biological differences between HPV+ and HPV-HNSCC, offering insights into HPV-driven oncogenesis. Understanding these distinctions may improve patient stratification and inform targeted therapeutic strategies for HNSCC.

## Introduction

Viruses are common carcinogenic agents, and one such virus is the Human Papillomavirus (HPV). HPV is a non-enveloped virus with a circular double-stranded DNA genome and infects basal cells of the stratified epithelium [1]. HPV infections are one of the most common sexually transmitted diseases, infecting millions of people every year [2]. Though most HPV infections are cleared spontaneously by the immune system, persistent infection with a high-risk HPV (type 16, 18, etc.) could lead to anogenital (e.g. cervical cancer, anal cancer, vulvar cancer, penile cancer) or head and neck squamous cell carcinoma (HNSCC) [3].

HNSCC is a collective term for the heterogeneous group of cancers of the mucosal epithelium of the oral cavity, paranasal sinuses, pharynx, oropharynx, lips, and larynx. These malignancies can be caused by carcinogens such as tobacco, areca nut, betel quid, and alcohol consumption, or from persistent infection with viruses such as Epstein-Barr virus (EBV) or HPV, especially HPV-16, and HPV-18[4-6]. In this article, we will focus on HPV and will therefore divide the HNSCC patients into two groups: HPV-positive (HPV+) or HPV-negative (HPV-). HPV-HNSCC cases are more prevalent in Southeast Asia and Australia, whereas HPV+ HNSCC cases are prominent in the USA and Western Europe. It has been observed that HPV+ HNSCC patients have a better prognosis, and the tumors are more sensitive to radiation and chemotherapy than HPV-HNSCC tumors. Additionally, men are more prone to develop HPV-HNSCC than women due to the aforementioned risk behaviors [7-12]. These data suggest fundamental differences in the cell biology of HPV-HNSCC and HPV+ HNSCC tumors. Therefore, it is essential to understand these differences for improving patient stratification and treatment approaches.

In this report, we analyzed the bulk RNA-seq data from the PanCancer Atlas 2018 dataset [13] using dimensionality reduction techniques such as Principal Component Analysis (PCA) and Uniform Manifold Approximation and Projection (UMAP) to study the differences between the HPV-HNSCC and the HPV+ HNSCC patients. Our analysis revealed that the HPV+ HNSCC cases formed a distinct cluster from the HPV-HNSCC cases. This clear clustering suggests that these cases are characterized by a specific gene expression signature, potentially reflecting unique molecular mechanisms associated with HPV-tumors. Upon further analysis, we observed that the cluster with HPV+ cases had upregulated expression of genes involved in cell cycle regulation and nucleic acid processing, including chromosome segregation and RNA splicing. In contrast, genes involved in apoptosis, skin, and epidermis development, among others, were downregulated as compared to clusters with HPV-cases. These findings highlight the molecular differences between HPV-positive and HPV-negative cancers and provide insights into the biological processes underlying HPV-driven oncogenesis.

## Method

### Data for the study

For analysis, we used the RSEM (Batch normalized from Illumina HiSeq_RNASeqV2) file and clinical data file from the HNSC PanCancer Atlas 2018 downloaded from the cBioPortal[14]: https://www.cbioportal.org/study/summary?id=hnsc_tcga_pan_can_atlas_2018

### Processing and analysis of the data

The data obtained was cleaned by removing duplicate rows and rows with missing or empty values. Then, the cleaned dataset was processed using the Seurat package v5[15]. Briefly, the Seurat object was created, followed by the normalization of the values. Variable features were obtained, and the data were scaled. Then, PCA was performed, followed by plotting an elbow plot to determine the principal components to use for the UMAP algorithm. The first 10 principal components were then used for the UMAP algorithm. Then, ggplot2[16] was used to generate and visualize the UMAP.

The patient clinical metadata was then merged with the metadata of the Seurat object, and these data were visualized using ggplot2.

### Gene ontology analysis

Gene markers that define a cluster via differential gene expression were identified for each cluster. These are a set of genes that are differentially expressed in a single cluster compared to all other cells. Then, for each cluster, upregulated (avg_log2FC > 0.25) or downregulated genes (avg_log2FC < -0.25) were selected. Enrichment analysis was performed using the enrichGO function from the clusterProfiler package [17]utilizing the biological process ontology. The top 5 upregulated and downregulated biological processes for each cluster were plotted and visualized using ggplot2.

## Results

### HPV+ HNSCC patients cluster together

To explore the gene expression profiles of the HPV- and HPV+ HNSCC, we performed dimensionality reduction techniques on the publicly available RNA-seq data from the PanCancer Atlas 2018 dataset (Fig. 1A). We only included the patients for whom most of the clinical data was available (Total: 515 patients, HPV-HNSCC: 417, HPV+ HNSCC: 96, HPV status indeterminate: 2). We chose to evaluate the gene expression profiles by PCA and UMAP, as these techniques help extract patterns and behavior from a complex biological dataset.

**Figure 1.**
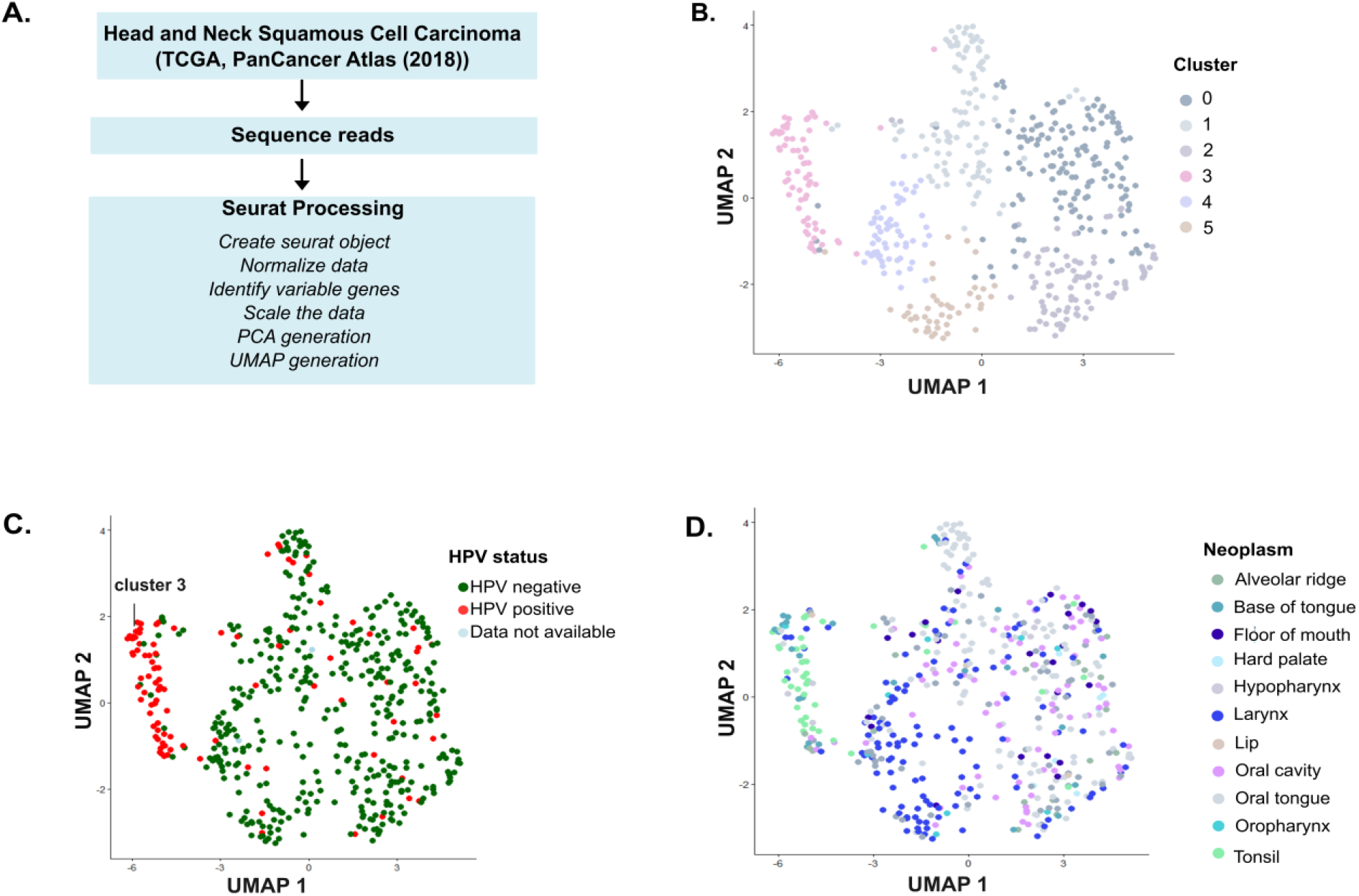
HPV+ HNSCC patients form a separate functional cluster from HPV-HNSCC patients. **(A)** Schematic showing the methodology used for the generation of PCA and UMAP plots. **(B)** UMAP plot showing the distribution of HNSCC patients in seven different clusters (0-5) based on their gene expression profiles. **(C)** The UMAP plot shown in (B) is colored based on the HPV status of the patients. 59.3% of HPV+ HNSCC patients were observed to be part of cluster 3. **(D)** The UMAP plot shown in (B) is colored based on the region of the patient’s neoplasm.

First, we employed PCA analysis, which transforms the original gene expression data into a number of principal components (PCs) that capture the most information in the data by finding linear combinations of the original features (in this case, genes) that best explain the variance in the dataset. Each PC represents the data that accounts for the highest variance. The first PC explains the highest variance, followed by the second PC, and so on. In our analysis, we observed that most of the HPV+ HNSCC patients clustered together (Fig. S1) when the first two PCs were plotted. This indicated that HPV-driven tumors exhibit a distinct molecular pattern compared to HPV-negative tumors.

Then, we wanted to see if the same trend is replicated with UMAP analysis, as UMAP analysis is used to investigate the structure of data in a lower-dimensional space, offering a different approach than PCA. While PCA seeks variance in a linear space, UMAP retains the local proximity of points while embedding the data in a 2D or 3D space. In this analysis, we observed that HNSCC patients formed 6 different clusters (clusters 0-5) (Fig. 1B), and while most of the HPV-HNSCC patients were present in different clusters, almost 59.3% of HPV+ HNSCC patients (57 out of 96 HPV+ HNSCC) were part of cluster 3 (Fig. 1C). This distinct clustering pattern of HPV+ HNSCC patients indicate that they have a common gene expression profile that the heterogenous expression profiles of HPV-HNSCC patients.

We also observed that most of the patients in cluster 3 had neoplasms in the base of the tongue, oral cavity, tonsil, or the hypopharynx (Fig. 1D).

### Gene Ontology (GO) enrichment analysis highlights distinct biological processes between HPV- and HPV+ HNSCC patients

Following the dimensionality reduction analysis, we wanted to know the biological significance of the clustering observed. For this, we used GO analysis to find out which groups of genes were upregulated or downregulated in clusters with 59.3% HPV+ HNSCC patients (cluster 3) (Fig. 2) and clusters with mostly HPV-HNSCC patients (cluster 0-2, 4, and 5) (Fig. S2 and S3).

**Figure 2.**
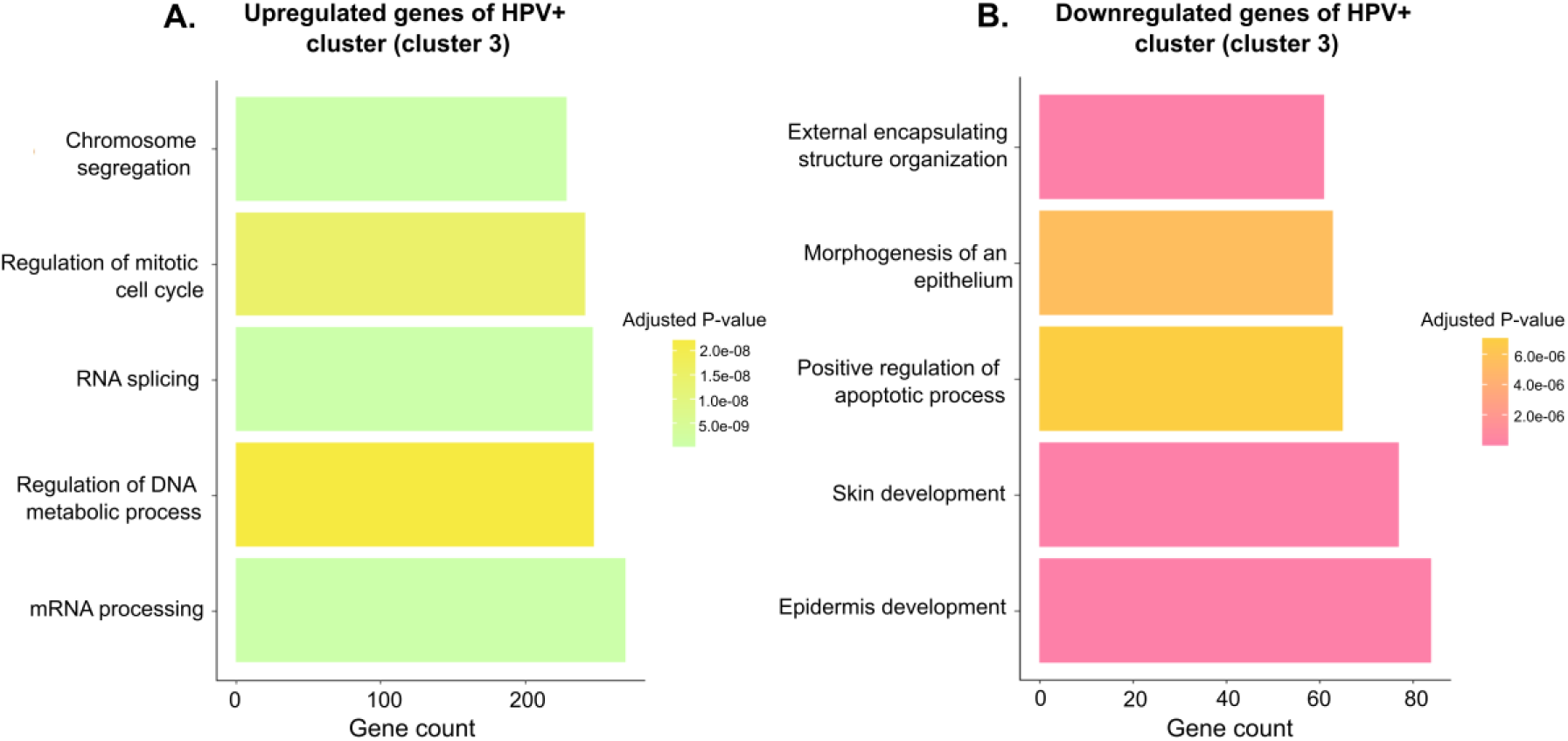
Genes supporting the replication of HPV are differentially regulated in HPV+ HNSCC patients. **(A)** Plot showing the upregulated biological process and the corresponding mRNA transcript counts for cluster 3. **(B)** Plot showing the downregulated biological process and the corresponding mRNA transcript counts for cluster 3.

We observed that in cluster 3 genes associated with nucleic acid processing such as mRNA processing, chromosome segregation, RNA splicing, etc. are upregulated (Fig. 2A). In contrast, genes associated with apoptosis, external encapsulating structure organization, epidermal development and skin development etc. were all downregulated (Fig. 2B). On the other hand, in clusters with the majority of HPV-HNSCC patients different sets of genes were upregulated or downregulated. Some genes were upregulated in one cluster and downregulated in another cluster, for instance, genes associated with blood circulation were downregulated in cluster 0 (Fig. S2A) but were upregulated in cluster 1 (Fig.S3B).

These observations suggest that in HNSCC several biological processes are altered, with the specific processes being contingent upon the presence or absence of HPV. Moreover, these results also highlight the heterogeneity of HPV-HNSCC patients.

## Discussion

HNSCC is the sixth most prevalent cancer globally with a high mortality rate. Its incidence is projected to increase by 30% by 2030 [18, 19]. A small percentage of the tumors of the oral cavity and the larynx and more than 70% of oropharyngeal cancer are associated with HPV [12].

This study analyzed the RNA-seq data from the PanCancer Atlas 2018 dataset using PCA and UMAP analysis. In our study, we observed clear clustering of the majority of HPV+ HNSCC patients (Fig. 1C), reinforcing the hypothesis that HPV+ HNSCC is a disease by itself and follows a distinct molecular trajectory from HPV-HNSCC, which is more heterogeneous in its gene expression profiles (cluster 0-2,4-6, Fig. 1B).

A small percentage of HPV+ HNSCC patients clustered together with HPV-HNSCC patients outside cluster 3. This could be due to inadequate viral load, which could significantly change their overall gene expression. It is also possible that these patients had an HPV infection and developed HNSCC from an environmental exposure or other cause, independent of the infection. However, the fact that these patients show a unique gene expression profile highlights the possibility that some HPV+ cancers may behave more like HPV-cases.

Moreover, it shows the potential utility of UMAP analyses for identifying unique cancer subtypes. Overall, these data suggest that HPV infection plays a central role in shaping the tumor transcriptome and underscores the importance for HPV+ HNSCC to be treated differently from HPV-HNSCC [20].

The HPV DNA encodes two types of protein: late proteins (L1 and L2, which form the viral capsid) and early proteins (E1-E8, which help the viral DNA replication, transcription, and host cell transformation). The expression of these proteins is tightly regulated through multiple HPV genome-encoded promoters, RNA splicing, and epigenetic modifications. HPV infects basal cells, and during a productive infection, the viral gene expression is synchronized with the differentiation of the host cell. So, as the infected cell moves toward the surface, the viral genome is replicated and packaged in capsid proteins. These infectious virus particles are then released. But, in the case of high-risk HPV, this infection cycle is disrupted as these types encode highly potent oncoproteins E5, E6, and E7. These proteins dysregulate several cellular processes, making the cells prone to oncogenic transformation. For instance, E6 targets the tumor protein 53 (p53) for degradation, which prevents apoptosis, whereas E7 targets retinoblastoma (Rb), a key regulator of cell cycle progression from the G1 to S phase. Degradation of Rb leads to unchecked continued cell cycle progression instead of differentiation.[1, 21, 22].

Our differential gene expression analysis revealed that genes associated with cell cycle and nucleic acid processing were upregulated, whereas genes related to epidermis development and morphogenesis were downregulated in HPV+ cases (Fig. 2). This suggests that HPV+ tumors maintain a gene expression profile that favors viral replication and proliferation rather than epithelial differentiation. This aligns with the activity of HPV oncoproteins, which drives continuous cell division at the expense of terminal differentiation. Also, the downregulation of epidermis and skin development in HPV+ cases is in agreement with a comprehensive study of routine clinical practice, involving 435 patients that showed that nonkeratinizing squamous cell carcinoma is strongly associated with high-risk HPV[23]. In addition to this, genes associated with apoptosis were downregulated, which reflects the involvement of the HPV oncoproteins described above.

Our differential gene expression analysis also reiterated the heterogeneity of HPV-HNSCC and revealed that genes corresponding to different biological processes were upregulated and downregulated in different HPV-HNSCC clusters (Fig. S2 and S3). The complexity of HNSCC explains why different patients respond differently to therapy [24].

Previous studies have used single-cell RNA sequencing to analyze the immune microenvironment of HPV+ and HPV-HNSCC subtypes, and HPV integration events in detail, and have shown that HPV+ HNSCC has very different genomic, mutational, transcriptional, and immunological profiles than HPV-HNSCC[25-27]. Our study builds upon these findings by leveraging bulk RNA sequencing data from the PanCancer Atlas 2018 dataset and examines the broader transcriptional landscape of HPV+ and HPV-HNSCC. While single-cell studies provide high-resolution insights into the individual cell level, our approach allows a practical assessment of variable gene expression levels across entire populations by analyzing bulk gene expression. We can identify overreaching trends, detect global regulatory mechanisms, and draw conclusions about the collective behavior of patient populations to better understand disease progression.

In summary, our study reinforces the notion that HPV+ HNSCC displays a divergent gene expression pattern and that these patients require tailored treatment. The unique gene expression pattern of HPV+ cases suggests potential differences in tumor biology, immune response, and treatment sensitivity, which could have significant clinical implications.

Future investigation is required to understand the precise mechanism by which HPV+ and HPV-HNSCC diverge at the molecular level. Understanding the drivers of these different pathways could provide valuable insights into tumor progression and identify potential biomarkers for early detection.

## Acknowledgements

We thank all colleagues in the Walker lab for their help and support, especially Kristen Dias, for their input on the bioinformatics workflow. The authors were supported by funding from the NIH (R01GM139977 to S.E.W)

