## Supplemental Data for "Analysis of Divergent Gene Expression between HPV+ and HPV- Head and Neck Squamous Cell Carcinoma Patients"

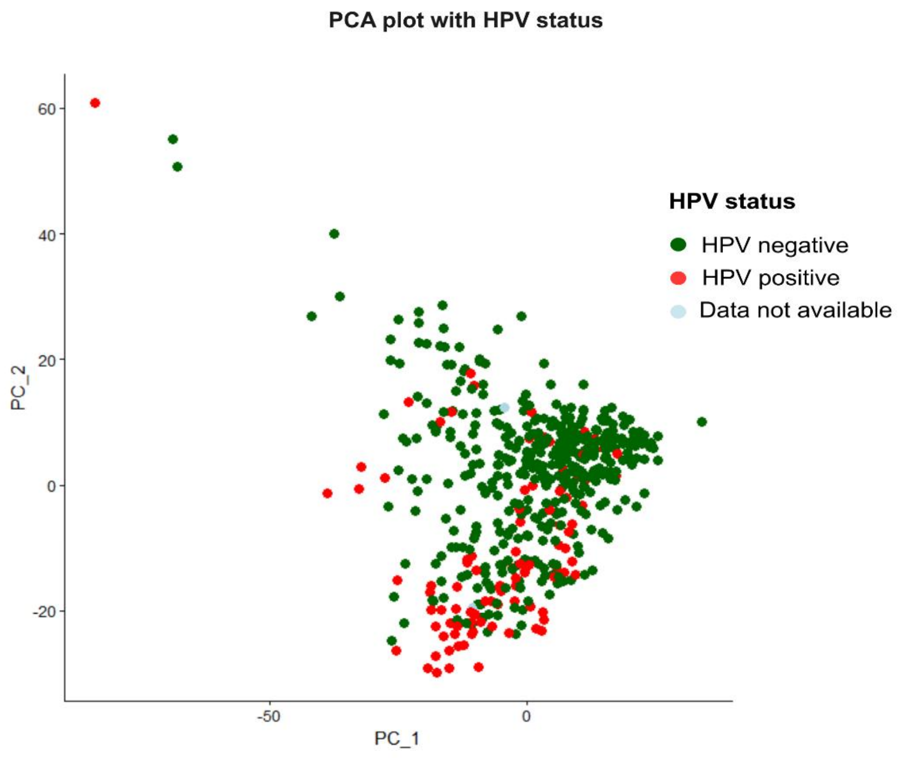


**Supplementary Fig. 1- Plot showing the distribution of HNSCC patients colored on their HPV status**

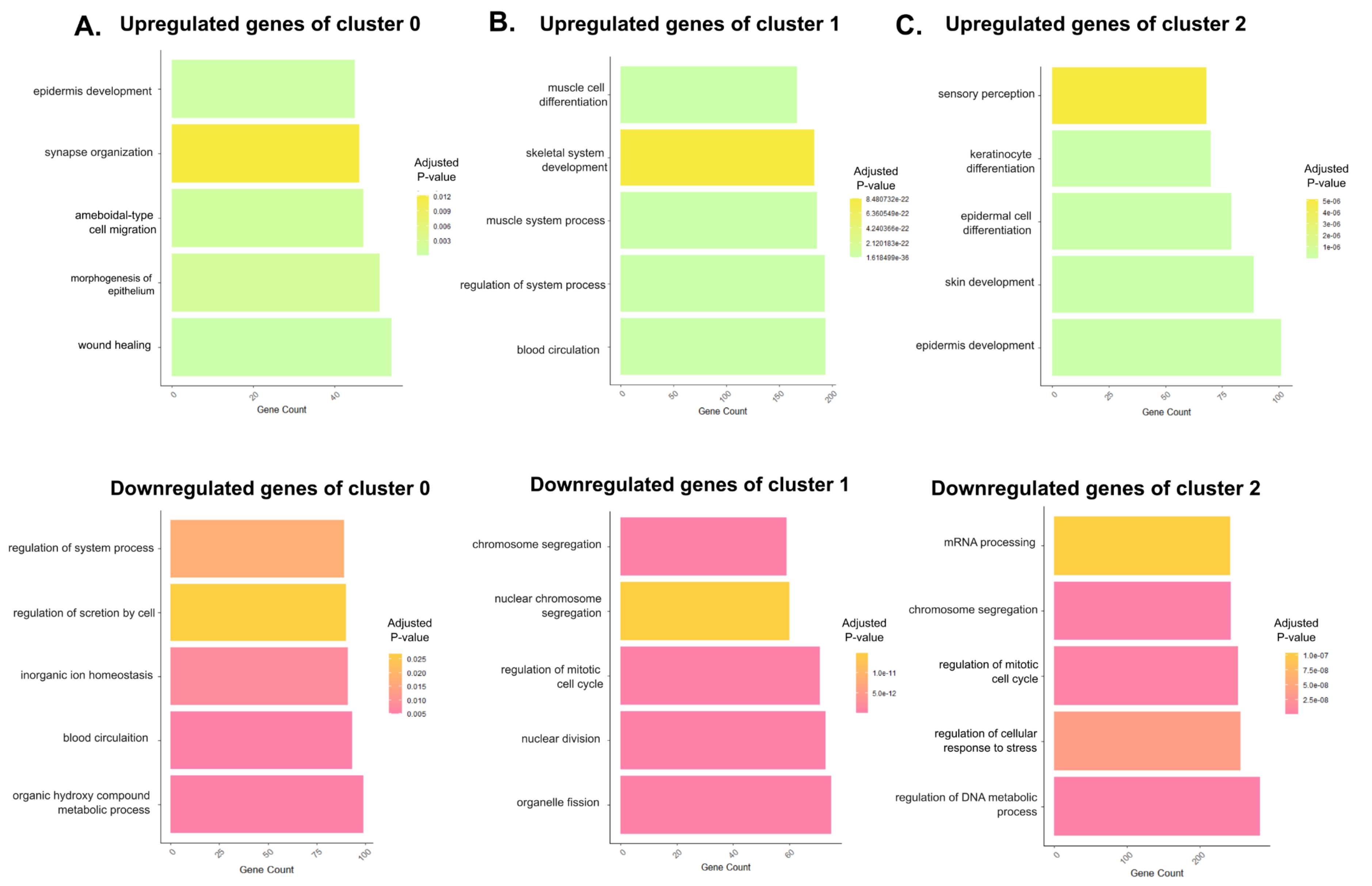


**Supplementary Fig. 2- Plot showing the upregulated and downregulated biological process and the corresponding mRNA transcript counts for clusters 0-2 (A-C)**

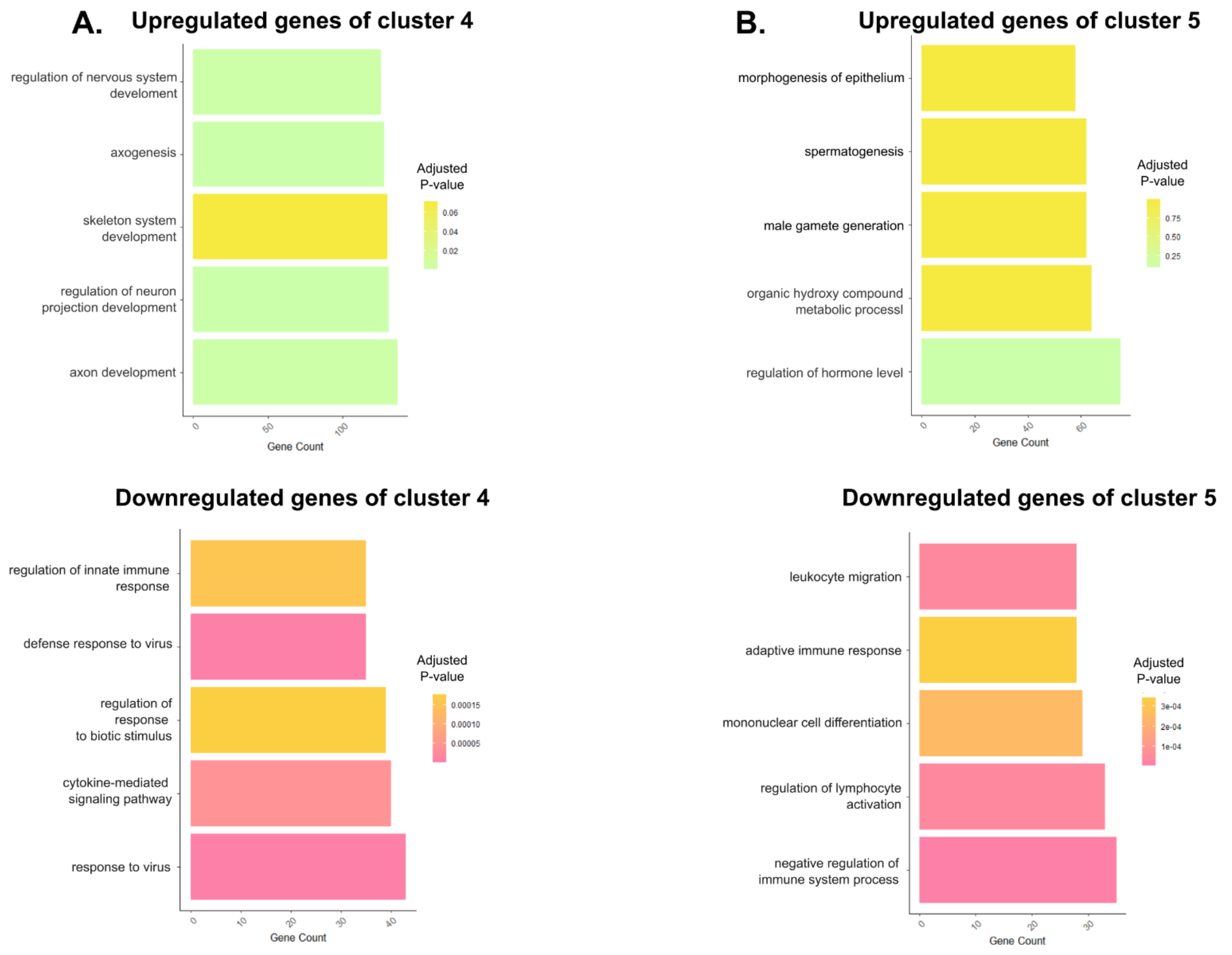


**Supplementary Fig. 3- Plot showing the upregulated and downregulated biological process and the corresponding mRNA transcript counts for clusters 4 and 5 (A and B)**
